# Experimental validation of graded arithmetic and linguistic working-memory tasks using behavioral and cerebral hemodynamic measures

**DOI:** 10.64898/2026.07.17.739087

**Authors:** Héctor Rojas-Pescio, Marco Villalta, Max Chacón

## Abstract

Graded cognitive tasks are relevant only when increasing nominal difficulty produces measurable changes in performance and a convergent physiological response. This study evaluated a five-level arithmetic paradigm and a five-level linguistic paradigm designed to engage working memory through separate exposure, retention, and response intervals. Thirty-eight healthy young adults completed both paradigms. Behavioral manipulation validity was assessed from accuracy and completion time; linguistic item-level responses were additionally modeled to detect nonlinear floor effects. Bilateral middle cerebral artery blood-flow velocity was recorded with transcranial Doppler ultrasonography. Baseline-normalized mean velocity change was the primary physiological endpoint, with positive and net area under the curve, peak response, time to peak, and early slope as secondary endpoints. Accuracy declined and completion time increased strongly across levels in both domains, with a substantially steeper accuracy decline in the linguistic paradigm. Although the levels were nominally ordered, the higher linguistic levels did not produce equidistant increases due to the ground effect. Transcranial Doppler models showed significant level-dependent changes in mean velocity, positive and net area under the curve, time to peak, and early slope. However, domain-by-level and domain-by-level-by-hemisphere interactions were not significant. Behavioral and physiological measures therefore converged mainly at the level of the graded experimental manipulation, rather than as distinct domain-specific or lateralized hemodynamic signatures. These findings lend support to paradigms as a multimodal framework for manipulating cognitive load experimentally, while also indicating upper-level language saturation and physiological quality-control decisions that should guide further improvement.

## 1. Introduction

Cognitive workload arises when the processing and temporary storage demands of a task approach the limited capacity of working memory (Sweller, 1988; Paas, Renkl & Sweller, 2003; Xie & Salvendy, 2000). The n-back family of paradigms has been used extensively to manipulate memory load, attention, updating, and response selection (Kirchner, 1958; Kane et al., 2007; Meule, 2017; Just & Carpenter, 1992). Nevertheless, nominal changes in task level do not by themselves establish valid graded manipulation. A defensible experimental paradigm should demonstrate that increasing difficulty produces systematic performance changes, avoids uninterpretable ceiling or floor effects, and elicits convergent physiological variation.

Arithmetic and linguistic tasks impose partially distinct processing requirements. Arithmetic operations require maintenance and manipulation of numeric information, whereas reading comprehension combines temporary storage with semantic integration, retrieval, and inference (Carpenter, 1992; Vingerhoets & Luppens 2001; Galy & Mélan, 2015). These differences make the two domains useful for investigating whether a graded manipulation generalizes across task content. At the same time, differences in item number, exposure duration, response format, and baseline ability can generate nonlinear performance trajectories. Validation must therefore distinguish a general level effect from domain-specific effects and from saturation at the most difficult levels.

The graded task battery was a study-specific experimental instrument designed to manipulate cognitive demand rather than a standardized psychometric test. Its validity evidence was based on content alignment with working-memory (Miyake & Shah, 1997) and cognitive-load theory (Paas, Renkl & Sweller, 2003), expert review of the arithmetic and linguistic items, pilot testing to avoid excessive frustration or fatigue, software-controlled administration, and empirical manipulation checks based on accuracy, completion time, and TCD endpoint trajectories. Because the task levels were intentionally constructed to differ in difficulty, internal consistency indices such as Cronbach’s alpha were not considered appropriate evidence of reliability. Instead, procedural reliability was addressed through standardized timing, fixed exposure-retention-response phases, computerized response recording, and uniform administration across participants.

Transcranial Doppler ultrasonography provides continuous, non-invasive measurement of blood-flow velocity in the middle cerebral arteries (MCA) and has been used to characterize neurovascular responses during cognitive activation (Moody, Panerai, et al., 2005; Panerai, et al., 2012; Csipo, et al., 2021; Intharakham, et al., 2022; Beishon, et al., 2017; Beishon, et al., 2018a; Beishon, et al., 2018b; Panerai, et al., 2019). Previous studies have shown that cognitive paradigms can produce bilateral flow-velocity changes, but the sensitivity of conventional TCD endpoints to increasing task complexity is not uniform. Some graded working-memory studies have reported larger responses under higher workload in both arithmetic and linguistic tasks (Badcock & Groen, 2017; Csipo, et al., 2021), whereas other paradigms have not yielded a reliably scalable conventional flow-velocity metric (Intharakham, et al., 2022). These observations motivate an integrated validation strategy rather than reliance on a single physiological measure.

The present study proposes a multimodal validation framework (Figure 1) that reframes a two-domain experimental protocol as a multimodal validation of graded cognitive manipulation. This multimodal strategy is consistent with current recommendations that cognitive workload should be assessed through convergent behavioral, subjective, and physiological indicators rather than inferred from a single measure (Charles & Nixon, 2019). The primary objectives were to determine whether five arithmetic and five linguistic working-memory levels produced (i) progressive behavioral change, (ii) level-dependent cerebral hemodynamic change, and (iii) convergence between behavioral and physiological measures. Domain-specific and hemispheric effects were evaluated as secondary questions. Subjective workload and information-theory measures were treated as supporting sensitivity analyses rather than primary validation evidence.

**Figure 1.**
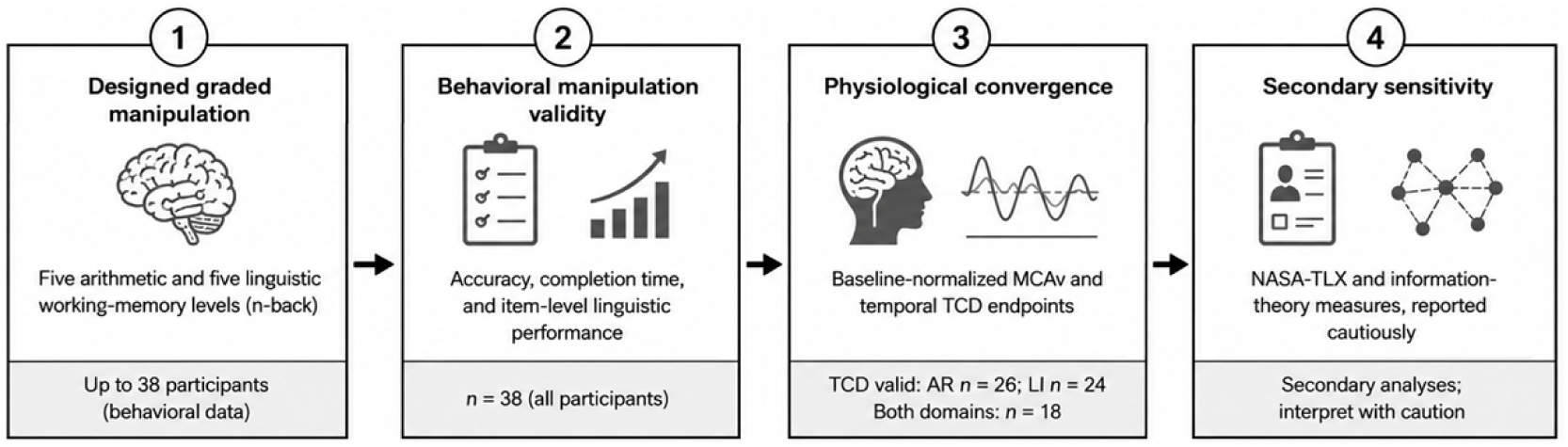
Multimodal validation framework used to organize the revised manuscript. Behavioral manipulation validity and conventional TCD endpoints constitute the primary evidence; subjective workload and information-theory measures are secondary.

## 2. Materials and Methods

### 2.1 Subjects and measurement

This study involved 38 healthy young adult volunteers (mean age: 25.21 ± 1.94 years; 55% female; 90% right-handed), recruited from the Universidad de Santiago de Chile. Six participants were excluded due to excessive motion artifacts or poor-quality blood pressure signals. Participants reported no history of cardiovascular, neurological, or psychiatric conditions and were not on medication at the time of testing. All subjects provided written informed consent in accordance with the Declaration of Helsinki. The study protocol was approved by the University Ethics Committee (Approval No. 280, dated May 8, 2023).

To assess cognitive effort, participants performed graded arithmetic (AR) and linguistic (LI) working-memory tasks while bilateral cerebral blood velocity (CBv) was recorded using transcranial Doppler ultrasound (TCD). CBv was measured in the left and right MCA using a DWL Doppler-Box system with two 2 MHz transducers mounted on an adjustable headframe to ensure probe stability throughout the session.

Arterial blood pressure (BP) was continuously recorded using a Finapress MIDI device with a photoplethysmographic cuff on the non-dominant hand. End-tidal CO₂ was not available for the present analysis. Therefore, no adjustment for CO₂ was performed, and the observed MCAv changes cannot be interpreted as CO₂-controlled neurovascular responses (Tarantini et al., 2019). However, environmental conditions were monitored and standardized across sessions to reduce variability. The within-subject task design and standardized task blocks aimed to minimize systemic drift in respiratory parameters, and the primary physiological endpoint was baseline-normalized mean MCA velocity change as shown in eq. (1).

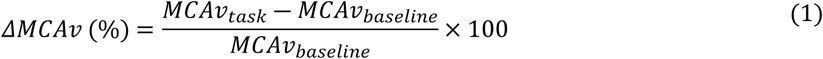

The endpoint was calculated per participant, domain, level, and hemisphere. Secondary temporal endpoints included peak percent change, net and positive area under the curve (AUC), time to peak, slope to peak, first-30-s slope, final-10-s residual, and peak-to-final change. The final-10-s measure represents the tail of the task recording and not a separate post-task recovery period.

All participants were instructed to maintain a normal and constant level of respiration throughout the duration of the task, which was completed in a quiet, temperature-controlled room with consistent lighting. To sustain engagement and motivation, participants received a financial stipend (via ANID doctoral research project funds) and were informed of an additional incentive awarded to the highest-performing participant, based on accuracy and perceived cognitive effort.

CBv and BP signals were sampled at 100 Hz and stored in standard PAR file format using QL Software v3.5.8 (DWL, Germany). Timing synchronization between the stimulus presentation software and the TCD recording system was achieved by aligning system clocks via the host operating system and validated NTP protocol, ensuring approximate temporal alignment for event markers across cognitive task levels. CBv and BP signals were exported in PAR format and preprocessed using MATLAB (MathWorks, R2023a). To obtain standardized beat-to-beat hemodynamic signals for endpoint extraction, the CBv signals underwent artifact removal and smoothing using a combination of Spline interpolation, Hampel filtering, and a fourth-order Butterworth low-pass filter (cutoff: 0.5 Hz) to suppress cardiac and respiratory noise components while preserving cognitive task-related low-frequency variations.

Beat detection was performed to identify systolic peaks and compute beat-to-beat mean values of BP and CBv. These were then interpolated using third-order polynomials and resampled at a constant rate of 5 Hz, following standard procedures for cerebral hemodynamic analysis (Moody, et al., 2005). The final signals for each domain were segmented per task level using synchronized timestamps, resulting in six time-series segments per participant (baseline plus five task levels).

Before each session, the experimental environment was checked for ambient noise, and participants received standardized instructions to reduce movement and maintain a steady breathing rate during recordings. Signal quality was visually verified before data collection began. According to the TCD signals acquisition protocol, participants were excluded if their CBv or BP signals exhibited >20% of continuous signal dropout or non-physiological amplitude spikes lasting more than 5 seconds, despite artifact correction attempts. Consequently, recordings from six participants were excluded due to excessive motion artifacts or poor-quality blood pressure signals caused by sweating or transient cuff displacements, resulting in a final valid sample of 38 participants for behavioral analysis, 26 participants for arithmetic tasks, 24 participants for linguistic tasks, and 18 participants for both domains.

### 2.2 Graded arithmetic and linguistic paradigms

The paradigms were n-back-inspired working-memory tasks rather than canonical identity-matching n-back trials. Each task separated stimulus exposure, a retention interval, and a response interval. Levels were presented in sequential experimental blocks, and a 90-s rest period followed each level as presented in Figure 2. The software standardized task presentation and recorded level accuracy and completion time. This approach is commonly used to elicit working memory and attentional demands in cognitive load paradigms (Kirchner, 1958; Sweller, 2010; Paas et al., 2003). The design aimed to progressively increase task difficulty while controlling time-on-task and participant fatigue.

**Figure 2.**
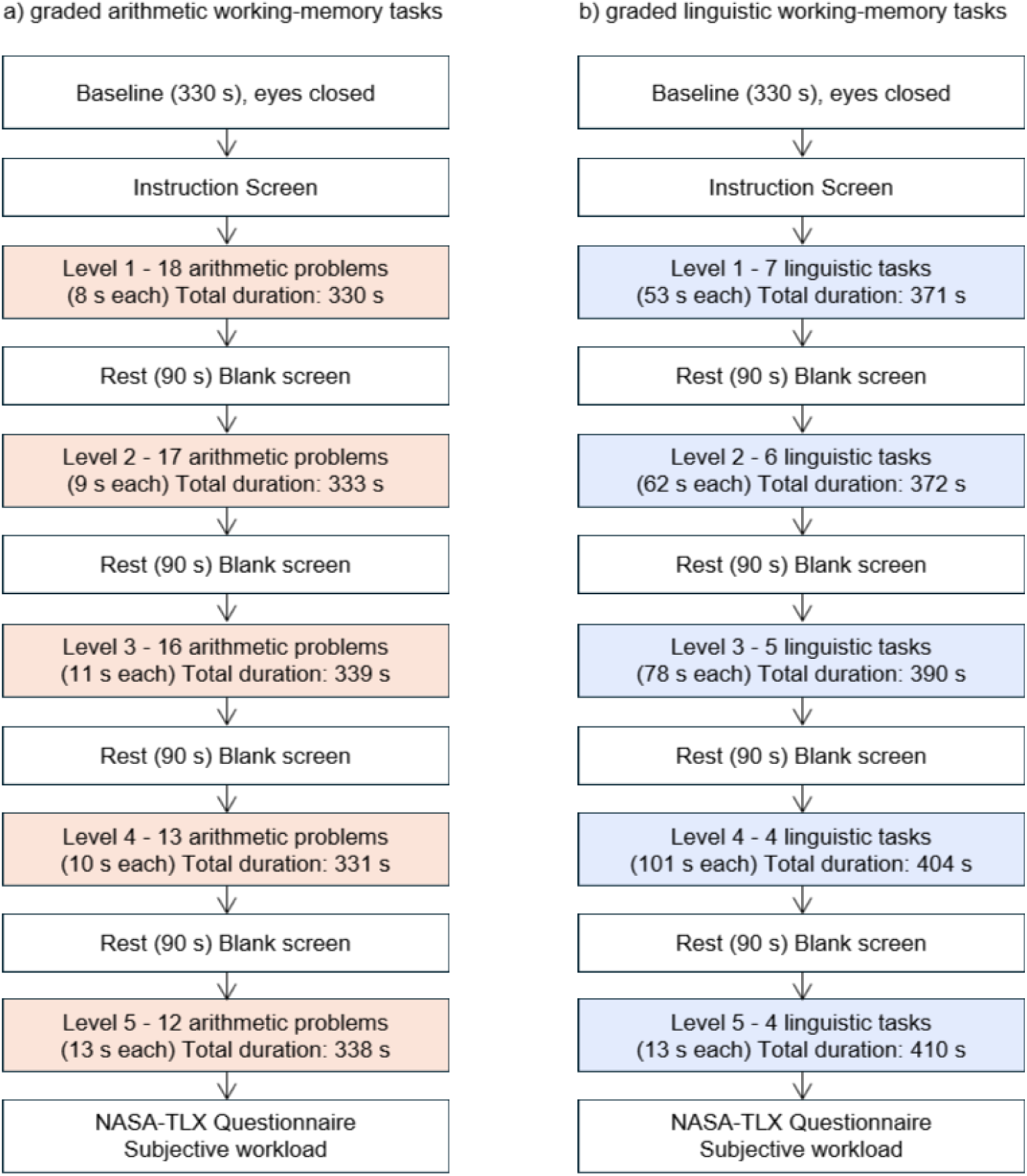
Experimental protocol for the graded working-memory paradigms. (a) Arithmetic paradigm comprising five ordered levels of increasing numerical processing and retention demands. (b) Linguistic paradigm comprising five ordered levels of increasing textual, semantic, and retention demands. Consecutive levels were separated by 90-s resting periods. Behavioral accuracy and completion time were recorded throughout both paradigms, cerebral blood-flow velocity was acquired using bilateral transcranial Doppler monitoring, and subjective workload was assessed after completion of each domain using NASA-TLX. Detailed task configurations are provided in Appendix A1.

Arithmetic difficulty was manipulated through number of digits, carrying demands, and sequential additions, consistent with prior arithmetic workload paradigms that vary task complexity to modulate mental demand and performance (Galy & Mélan, 2015). Linguistic difficulty was manipulated by using reading-comprehension items with multiple-choice responses. Its designed score combined topic complexity, Webb cognitive complexity, integration cost, storage cost, terminology, writing level, and passage length.

#### 2.2.1 Task Characteristics

Two custom graded working-memory paradigms were administered: an arithmetic task and a linguistic task. Both paradigms comprised five ordered levels designed to progressively increase processing, and retention demands while maintaining approximately comparable task-block durations. The tasks were not conventional n-back tasks because participants were not required to compare the current stimulus with one presented (n) positions earlier. Instead, they were designed as graded working-memory tasks inspired by cognitive-load and retention paradigms.

In the arithmetic paradigm, participants mentally solved addition problems, retained the computed result during a brief stimulus-free interval, and subsequently entered their response using the keyboard. Pilot testing initially considered addition, subtraction, multiplication, and division; however, the final paradigm was restricted to addition reducing between-task heterogeneity, participant frustration, and dropout risk. Arithmetic difficulty was manipulated through three features: the number of digits in the operands, the presence or absence of carrying, and the number of sequential additions included in each task.

In the linguistic paradigm, participants read short passages, retained and integrated their content during a brief interval, and subsequently answered a multiple-choice comprehension question. Linguistic difficulty was manipulated using a study-specific index that incorporated topic complexity, Webb’s Depth of Knowledge, textual integration demands, number of key concepts to be retained, domain-specific terminology, writing complexity, and passage length. Because the linguistic index was developed specifically for the present experiment, it was used to construct ordered experimental conditions rather than as a validated psychometric scale.

The two paradigms were designed for university students from scientific and humanities-related disciplines. Pilot testing and expert feedback were used to ensure that the tasks required sustained cognitive effort without creating excessive frustration or fatigue. Nevertheless, subsequent behavioral analysis showed that the upper linguistic levels approached a performance floor. Therefore, the five levels are interpreted as ordered difficulty conditions rather than as psychometrically equidistant increments of cognitive effort.

Detailed difficulty definitions, task attributes, level configurations, timing parameters, and stimulus sequences are provided in Appendix A1, Design and Operationalization of the Graded Arithmetic and Linguistic Working-Memory Tasks.

#### 2.2.2 Experimental Protocol

Both paradigms were administered using a custom-built desktop application developed in the Microsoft .NET framework. The software controlled stimulus presentation, retention intervals, response windows, and progression through the task sequence. It also recorded behavioral responses, completion times, and timestamps for synchronization with physiological recordings.

Each arithmetic trial followed three consecutive phases:

1. **Stimulus exposure:** the arithmetic problem was displayed for a predefined interval.
2. **Retention interval:** the problem was removed from the screen, requiring the participant to retain and mentally process the result.
3. **Response window:** the participant entered the calculated answer within a fixed period.

Depending on task level, arithmetic exposure times ranged approximately from 5.2 to 14.0 s, retention intervals from 8.0 to 11.0 s, and response windows from 4.0 to 8.0 s.

Each linguistic trial followed a comparable three-phase structure:

1. **Stimulus exposure:** a passage was displayed for reading and comprehension.
2. **Retention interval:** the passage was removed, requiring the participant to retain and integrate the relevant information.
3. **Response window:** a multiple-choice question was presented, and the participant selected an answer.

Because linguistic passages differed in length and complexity, exposure times ranged from 30.0 to 66.0 s, retention intervals from 6.5 to 9.5 s, and response windows from 13.0 to 30.0 s.

The number of trials per level was adjusted according to task duration and complexity. Arithmetic levels contained between 12 and 18 trials, whereas linguistic levels contained between 4 and 7 trials. These adjustments maintained the total duration of each level at approximately 5.5 min. Mean level duration was approximately 334 s for the arithmetic paradigm and 331 s for the linguistic paradigm. It should be considered that comparability can be stated within each domain, but not between both.

A 90-s resting period separated consecutive levels to reduce cumulative fatigue and allow partial physiological recovery. Participants performed the tasks while sitting in front of a computer screen. Neither paradigm was self-paced: the software automatically advanced through the predefined exposure, retention, and response phases to ensure consistent timing across participants.

The five levels were administered in ascending order for all participants. Consequently, task level was partially confounded with elapsed time, habituation, and cumulative fatigue; this design feature was considered when interpreting level-dependent effects.

A baseline TCD recording with eyes closed of at least 330 s was obtained to provide a reference segment with a duration and number of observations comparable to those of the task levels. The same baseline recording was used for the current baseline-normalized analyses of both cognitive domains. Full timing configurations and the operational definitions of task difficulty are reported in Appendix A1.

Subjective workload was assessed after completion of each task domain using the NASA Task Load Index (NASA-TLX). Because NASA-TLX was collected at the domain level rather than after each individual level, it was used to compare overall subjective workload between arithmetic and linguistic tasks and not as a direct measure of the five-level gradient.

### 2.3 Statistical Analysis

Validation was organized into four evidence layers (Figure 1). First, behavioral manipulation validity required systematic level effects on accuracy and completion time. Participant-level linear slopes summarized ordered change, and repeated-measures models tested domain, level, and domain-by-level effects. Linguistic item-level correctness was analyzed with binomial GEE models including linear and quadratic level terms to test upper-level floor behavior (Appendix A2).

Second, physiological convergence was evaluated using Gaussian GEE models clustered by participant. Mean ΔMCAv was the primary endpoint; the temporal endpoints were secondary. Models included domain, centered task level, hemisphere, relevant interactions, and baseline MCAv. GEE was preferred for the unbalanced domain-specific datasets, while repeated measures ANOVA were used as a supplementary sensitivity analysis in the balanced n=18 subset. Greenhouse–Geisser was used as the primary correction and Huynh–Feldt as a supplementary check. Domain effects with (df=1) do not require correction. Two-sided p<0.05 was considered statistically significant.

Third, behavior-physiology convergence was explored using Spearman correlations across participant-domain-level observations and participant-level slopes. Multiple tests were controlled with the Benjamini-Hochberg false discovery rate (Benjamini & Hochberg, 1995). Behavior-adjusted GEE models tested whether accuracy or log completion time explained physiological variation beyond domain, level, hemisphere, and baseline MCAv. Fourth, domain-specific and hemispheric interactions were interpreted as secondary discriminant evidence rather than prerequisites for manipulation validity.

## 3. Results

### 3.1 Behavioral manipulation validity

Accuracy declined across levels in both domains, while completion time generally increased (Table 3; Figures 3-4). Arithmetic accuracy decreased from 98.38% at level 1 to 63.82% at level 5. Linguistic accuracy decreased from 98.12% to 32.24%, with most of the decline occurring by level 3. Arithmetic completion time increased from 297.00 s to 428.92 s, and linguistic completion time increased from 345.53 s to 483.34 s.

**Figure 3.**
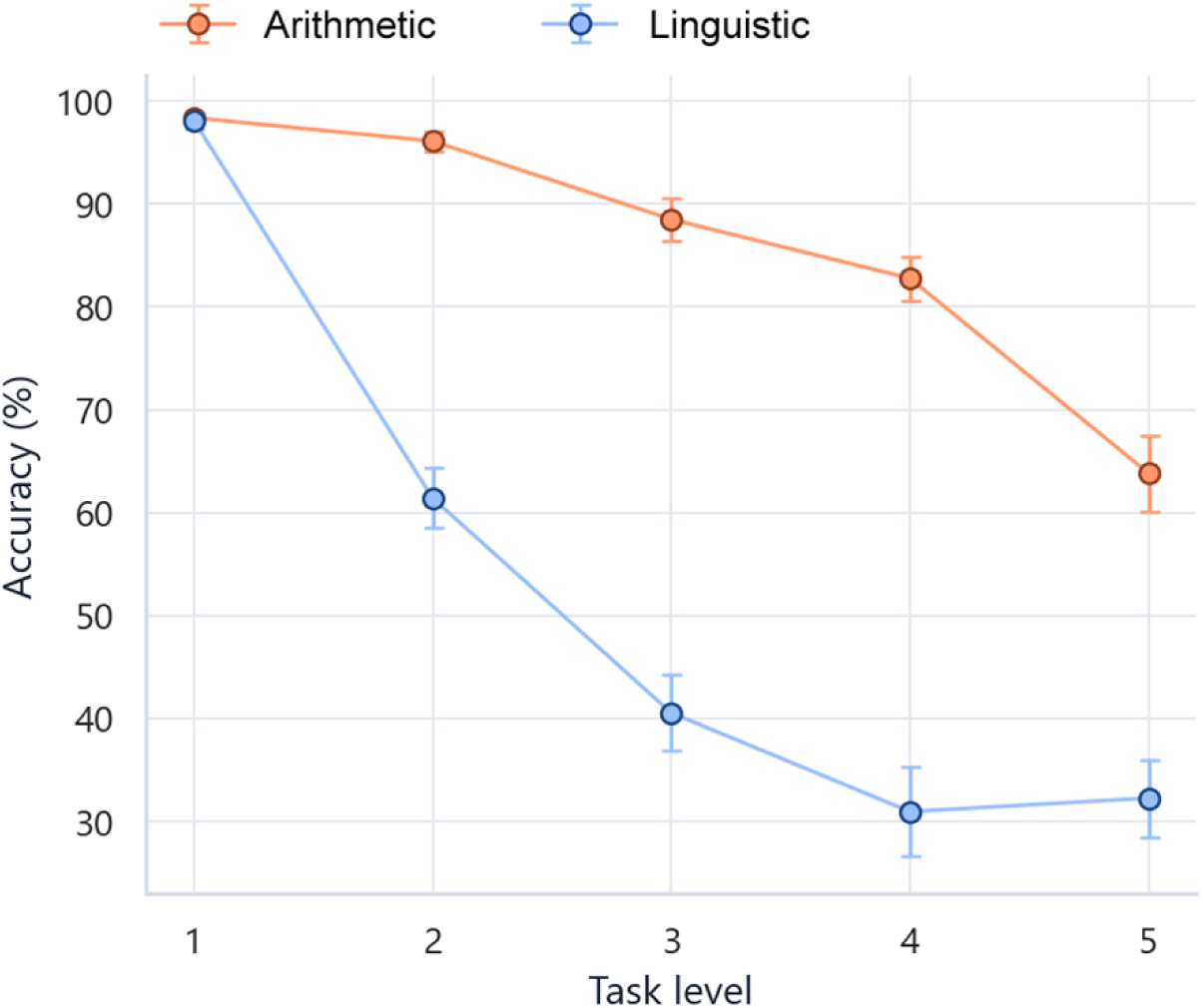
Mean accuracy by task level. Error bars represent standard errors across 38 participants.

**Figure 4.**
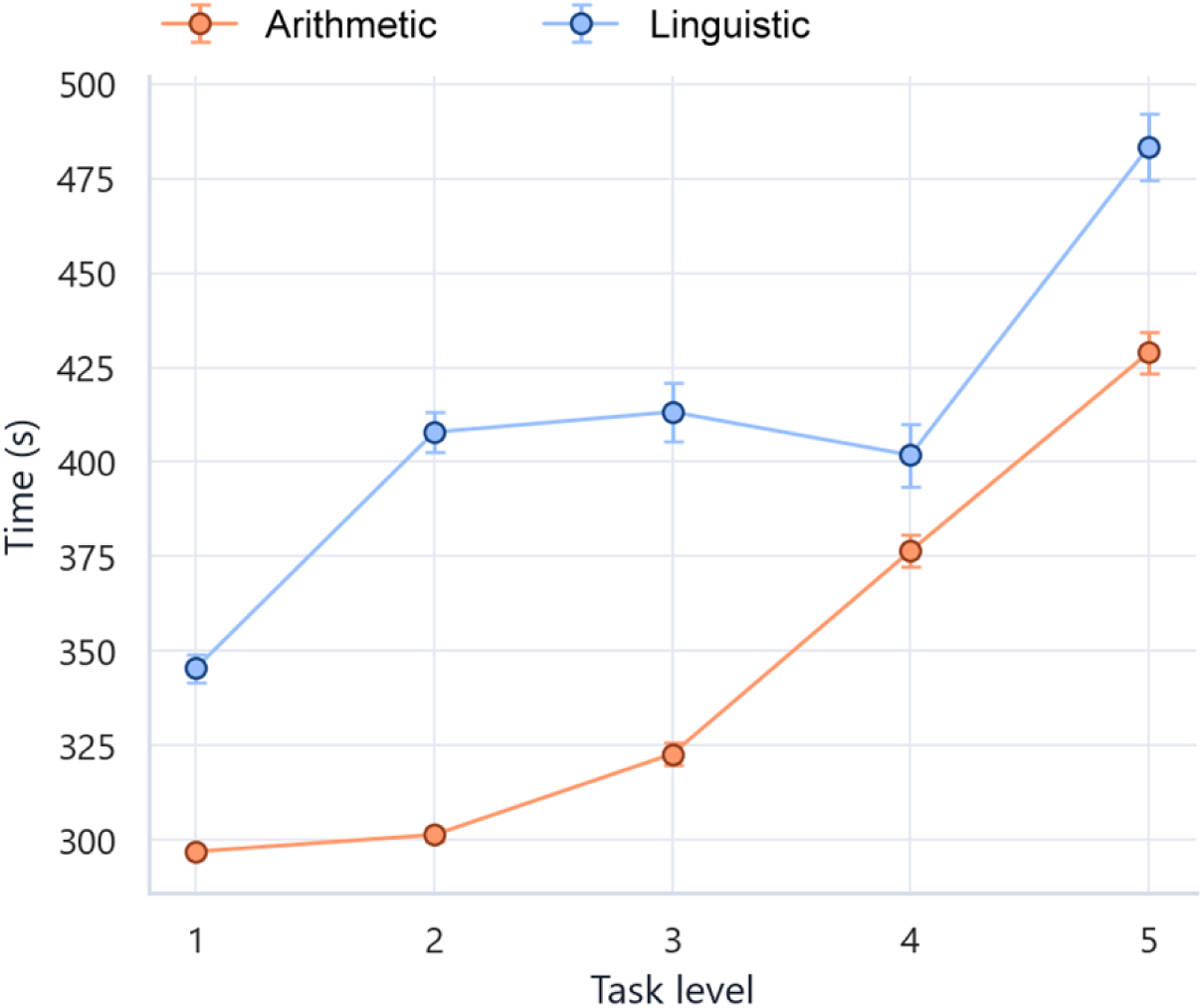
Mean completion time by task level. Error bars represent standard errors across 38 participants.

**Table 3.**
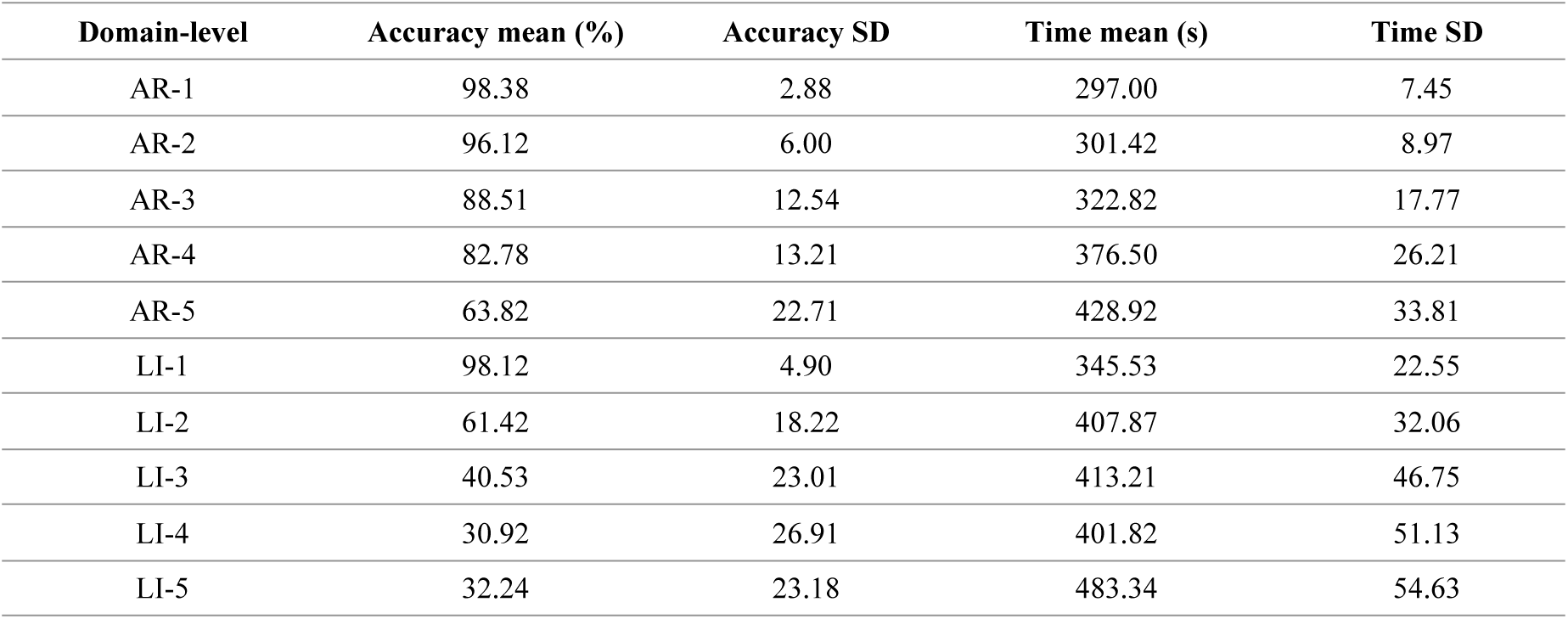
Behavioral performance by task domain and level (n=38).

| Domain-level | Accuracy mean (%) | Accuracy SD | Time mean (s) | Time SD |
| --- | --- | --- | --- | --- |
| AR-1 | 98.38 | 2.88 | 297.00 | 7.45 |
| AR-2 | 96.12 | 6.00 | 301.42 | 8.97 |
| AR-3 | 88.51 | 12.54 | 322.82 | 17.77 |
| AR-4 | 82.78 | 13.21 | 376.50 | 26.21 |
| AR-5 | 63.82 | 22.71 | 428.92 | 33.81 |
| LI-1 | 98.12 | 4.90 | 345.53 | 22.55 |
| LI-2 | 61.42 | 18.22 | 407.87 | 32.06 |
| LI-3 | 40.53 | 23.01 | 413.21 | 46.75 |
| LI-4 | 30.92 | 26.91 | 401.82 | 51.13 |
| LI-5 | 32.24 | 23.18 | 483.34 | 54.63 |

Participant-level accuracy slopes were negative for arithmetic (mean −8.25 percentage points per level, SD 5.08; t=−10.0, p=4.50×10⁻¹²) and linguistic tasks (mean −16.2, SD 5.93; t=−16.9, p=6.00×10⁻¹⁹). The paired linguistic-minus-arithmetic slope difference was −7.98 percentage points per level (SD 7.26; t=−6.78, p=5.62×10⁻⁸), indicating a steeper decline in linguistic performance. Log-time slopes were positive in both domains (AR=0.0951, p=1.29×10⁻²⁸; LI=0.0645, p=4.84×10⁻²⁰).

**Table 4.** Repeated measures behavioral effects with Greenhouse Geisser’s epsilon (*ε*_GG_) and corrected p value (*p*_GG_) in the current inferential analysis.

| Outcome | Effect | F | $\epsilon_{GG}$ | $p_{GG}$ |
| --- | --- | --- | --- | --- |
| Accuracy | Domain | F(1,37)=277.8 | 1.0000 | $8.698 \times 10^{-19}$ |
| Accuracy | Level | F(4,148)=121.7 | 0.7416 | $1.376 \times 10^{-34}$ |
| Accuracy | Domain × level | F(4,148)=31.64 | 0.7441 | $9.245 \times 10^{-15}$ |
| Log time | Domain | F(1,37)=184.8 | 1.0000 | $5.827 \times 10^{-16}$ |
| Log time | Level | F(4,148)=660.5 | 0.6179 | $1.314 \times 10^{-58}$ |
| Log time | Domain × level | F(4,148)=70.94 | 0.6700 | $3.266 \times 10^{-23}$ |

### 3.2 Linguistic upper-level floor behavior

Linguistic item-level analysis comprised 988 observed item trials. A linear binomial GEE showed decreasing log-odds of a correct response with level (β=−0.815, SE=0.0574, p=1.13×10⁻⁴⁵). In the quadratic model, the negative linear term was larger (β=−2.68, SE=0.176, p=2.93×10⁻⁵²) and the quadratic term was positive (β=0.447, SE=0.0422, p=2.79×10⁻²⁶). Together with the observed accuracy plateau near 31-40% across levels 3-5, this indicates a floor region rather than five evenly spaced increments in achieved difficulty.

### 3.3 Conventional TCD endpoint trajectories across task levels, domains, and hemispheres

The primary GEE analysis contained 500 hemispheric observations from the valid domain-specific TCD samples. Mean ΔMCAv decreased by 1.084 percentage points per level (SE=0.2566, 95% CI −1.587 to −0.581, p=2.40×10⁻⁵). The balanced repeated-measures sensitivity analysis also showed a level effect, F(4, 68)=17.1, p=1.69×10⁻^6^.

Several temporal endpoints showed convergent level effects. Positive AUC decreased by 508.2 %·s per level, net AUC decreased by 874 %·s per level, time to peak increased by 28.14 s per level, and the early 30-s slope increased by 0.0577 %/s per level (Table 5; Figure A3). These changes indicate an altered temporal hemodynamic profile with increasing task level rather than a simple monotonic increase in mean activation.

**Table 5.**
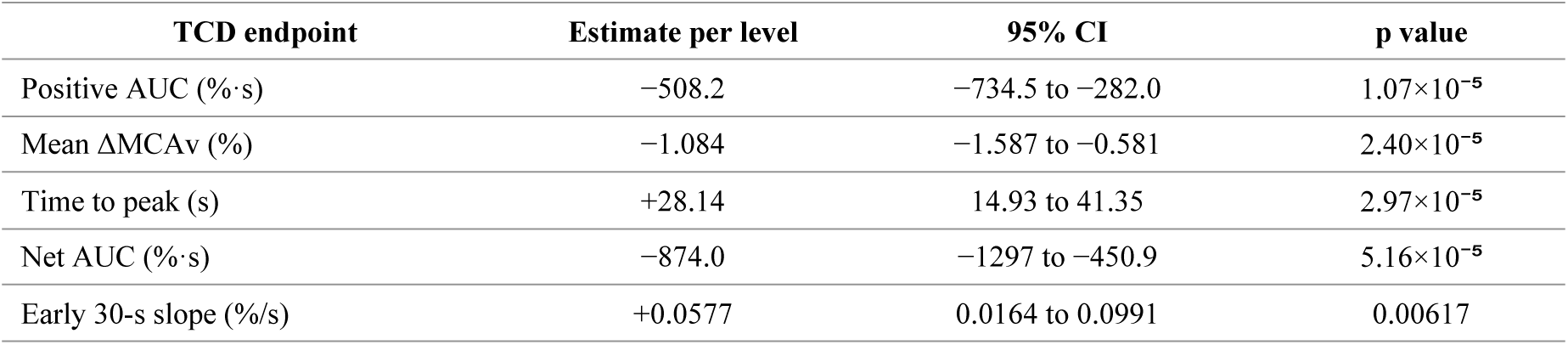
Strongest level terms from subject-clustered Gaussian GEE models of TCD endpoints.

The standardized endpoints with different units to be shown together; exact coefficients and confidence intervals can be seen in the Appendix A3 forest plot.

Figure 5 depicts domain- and hemisphere-specific trajectories of traditional TCD endpoints throughout task levels, with endpoint profiles emphasizing physiological changes at each level. The endpoint panel links the main claim to baseline-normalized MCAv while demonstrating if alternate temporal summaries give the same narrative. The displayed means are based on valid subjects (AR n=26, LI n=24), with a perfectly balanced cross-domain repeated-measures subset of n=18.

**Figure 5.**
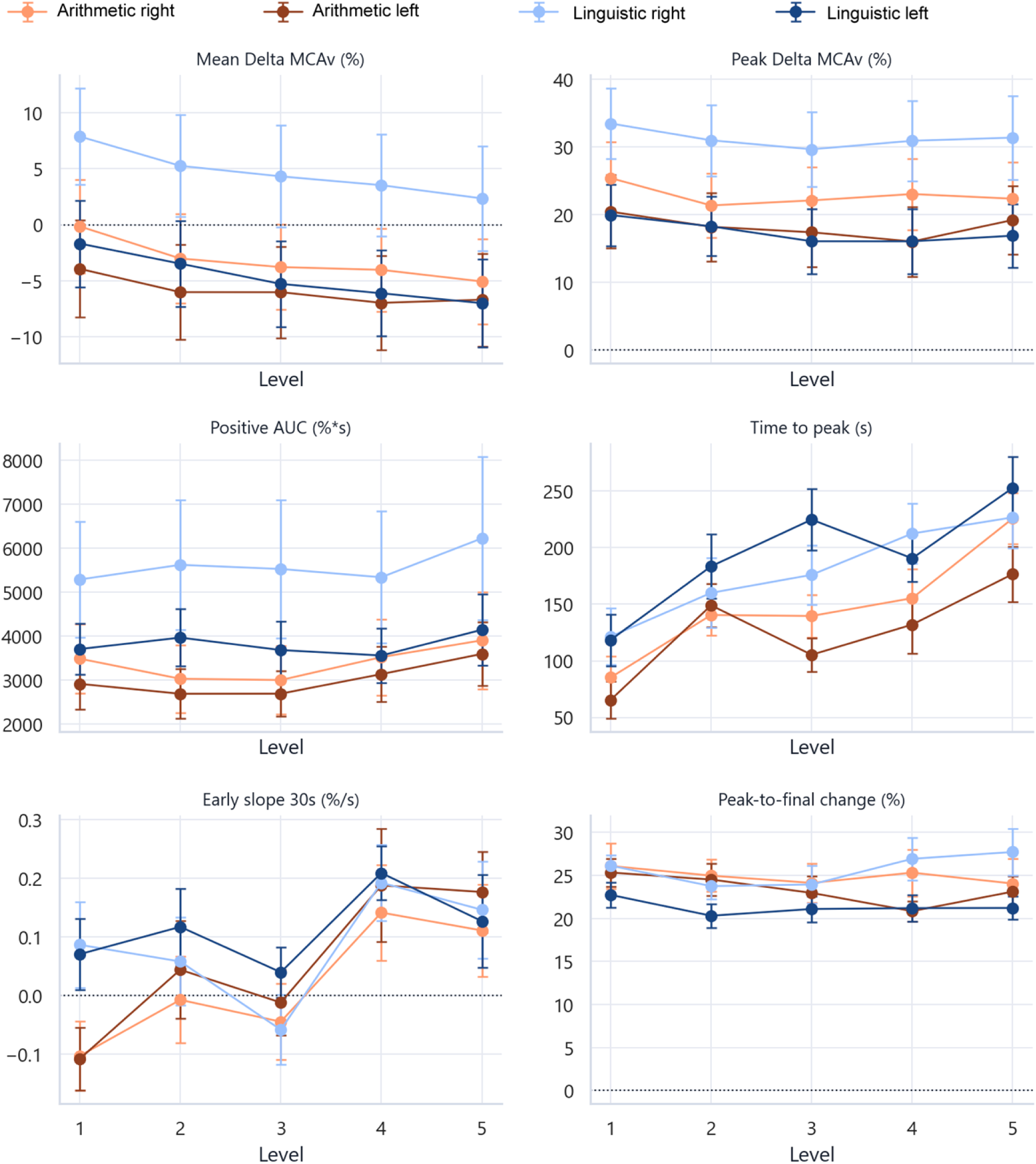
Domain- and hemisphere-specific trajectories of conventional TCD endpoints across task levels. Points represent mean values and error bars represent standard errors for participants meeting the final signal-quality criteria. Arithmetic and linguistic (right / left) denote right and left middle cerebral artery recordings. Mean ΔMCAv, peak ΔMCAv, positive AUC, time to peak, early 30-s slope, and peak-to-final change are shown by level. The trajectories illustrate the descriptive patterns underlying the population-averaged Gaussian GEE level effects reported in Table 5. Although several linguistic amplitude-related endpoints showed right–left separation, the omnibus hemisphere and domain-by-level-by-hemisphere effects were not statistically significant; hemispheric differences should therefore be interpreted as exploratory.

### 3.4 Domain and hemispheric specificity

The physiological level effect did not differ significantly by domain. In the GEE model, the linguistic-by-level term for mean ΔMCAv was β=−0.193 (p=0.604). In the balanced subset, the domain-by-level effect was F (4, 68)=0.655 (p=0.580). The hemisphere main effect (p=0.281), level-by-hemisphere effect (p=0.693), and domain-by-level-by-hemisphere effect (p=0.424) were also nonsignificant. The GEE three-way interaction was likewise nonsignificant (p=0.302). Laterality indices were therefore interpreted as exploratory and did not support a robust domain-specific hemispheric signature.

### 3.5 Behavioral-physiological convergence

Across participant-domain-level observations, longer completion time was associated with later TCD peak timing (Spearman ρ=0.360, FDR q=6.60×10⁻⁸), and lower accuracy was associated with later peak timing (ρ=−0.340, q=2.38×10⁻⁷). Longer completion time was also associated with greater positive AUC (ρ=0.227, q=0.00138), while lower accuracy was associated with a lower early 30-s slope (ρ=−0.191, q=0.00863).

However, participant-level correlations between behavioral and physiological slopes did not survive FDR correction. In behavior-adjusted GEE models, log completion time remained associated with positive AUC (β≈1206 %·s per standardized log-time unit, p=0.00153, q=0.0111), whereas most accuracy and time terms for other physiological endpoints were nonsignificant. Thus, behavior and physiology converged mainly through their common relation to task level rather than through tight participant-specific coupling.

### 3.6 Secondary subjective and nonlinear analyses

NASA-TLX scores were available at the domain level only and could not test the five-level gradient. The current total score was higher for arithmetic than linguistic tasks (54.82 vs 43.50). No NASA-TLX association with TCD endpoints remained significant after FDR correction in the integrated analysis.

Precomputed information-theory measures showed numerous level-related trends across entropy families, particularly normality-based, Rényi, Tsallis, and dispersion entropy configurations. Because many parameter combinations were screened and the final parameter-selection and subject-code mapping rules remain unresolved, these results were treated as supplementary sensitivity evidence rather than primary validation endpoints.

Overall, the results support multimodal convergence at the level of the ordered experimental manipulation, but not a tight participant-specific behavioral–physiological coupling or a robust domain-specific hemodynamic signature.

## 4. Discussion

This study provides multimodal evidence that the experimental protocol manipulated cognitive demand across ordered arithmetic and linguistic task levels. The strongest evidence was behavioral: accuracy declined, completion time increased, and the domain-by-level interactions showed that the two paradigms had different performance trajectories. Conventional TCD endpoints also varied by level, supporting physiological convergence. The results therefore justify describing the protocol as a graded manipulation, while also showing why validation cannot be reduced to nominal task labels or to a single hemodynamic amplitude measure.

### 4.1 Behavioral evidence and linguistic saturation

The arithmetic paradigm produced a progressive decline from near-ceiling accuracy at levels 1-2 to 63.8% at level 5, accompanied by increasingly long completion times. This pattern is consistent with a graded increase in demands for numeric maintenance and manipulation. The linguistic paradigm produced a much steeper decline, with accuracy near one third by levels 4-5. The significant quadratic item-level term confirms that achieved difficulty flattened at the upper end. Consequently, the linguistic task set is graded in the broad sense that higher designed levels are more difficult, but levels 3-5 should not be treated as psychometrically equidistant. Once performance approached the floor at levels 3–5, additional nominal complexity may have increased frustration or disengagement without producing measurable behavioral discrimination. Under these conditions, hemodynamic changes may reflect sustained overload, reduced effort investment, or fatigue rather than progressively increasing successful cognitive processing.

For future applications, the upper linguistic levels should be recalibrated by reducing passage complexity, adjusting distractor quality, increasing the number of items, or applying adaptive branching. Such changes could preserve discrimination between participants while avoiding a floor that limits dose-response interpretation. The arithmetic paradigm may also benefit from additional item-level capture so that both domains can be analyzed with equivalent trial-level models.

The higher subjective workload reported for arithmetic tasks, despite markedly poorer linguistic accuracy, suggests that subjective effort and objective performance captured different aspects of task engagement. Participants may have remained actively engaged in the arithmetic problems because successful solutions appeared attainable, thereby reporting greater mental demand and effort. Conversely, the early performance collapse in the linguistic paradigm may have promoted disengagement or reduced effort investment at the upper levels, producing lower subjective workload despite poorer accuracy. Mathematics-related anxiety or culturally reinforced perceptions of arithmetic difficulty may also have contributed, although these mechanisms were not directly measured.

These findings have theoretical and empirical implications for cognitive-workload measurement. Behavioral performance captured the achieved difficulty of the task with high sensitivity, particularly through accuracy decline and completion time increase. However, behavioral measures were also domain-dependent and vulnerable to ceiling and floor effects, as shown by the upper-level saturation of the linguistic paradigm. In contrast, cerebral hemodynamic endpoints provided independent physiological evidence that ordered task levels modified task-related MCAv dynamics. Nevertheless, the absence of robust domain-by-level or domain-by-level-by-hemisphere effects suggests that conventional MCAv endpoints primarily reflected a broad task-demand response rather than a specific arithmetic, linguistic, or lateralized neurovascular signature. Therefore, behavioral and physiological measures should be interpreted as complementary rather than interchangeable indicators of cognitive effort.

### 4.2 Hemodynamic convergence is level-dependent but not domain-specific

The significant level term for mean ΔMCAv and the convergent effects in AUC, time to peak, and early slope indicate that task progression altered the cerebral hemodynamic response. Importantly, the direction was not a simple increase in mean flow velocity with greater designed difficulty. Positive and net AUC decreased, while peak timing was delayed. Prior fTCD studies likewise show that cognitive responses depend on temporal dynamics, systemic influences, task duration, and the form of stimulation (Moody, Panerai, et al., 2005; Panerai, et al., 2012; Csipo, et al., 2021; Intharakham, et al., 2022; Panerai, et al., 2017). The present pattern may reflect sustained effort, habituation, delayed recruitment, fatigue, or differences between the common baseline and task blocks; the available data do not distinguish these mechanisms.

Because all task levels were presented in ascending order, the effect of nominal difficulty cannot be fully separated from elapsed time, habituation, learning, disengagement, or cumulative fatigue. The progressive reduction in mean ΔMCAv and AUC may therefore represent an adaptation to repeated stimulation or declining task engagement, rather than a direct inverse dose-response relationship between cognitive demand and cerebral blood-flow velocity. The fixed sequence was selected in the original design to provide a controlled progression and reduce early frustration or withdrawal; however, future studies should counterbalance or randomize block order.

The absence of significant domain-by-level and three-way hemisphere interactions limits claims of domain-specific neurovascular signatures. This does not undermine manipulation of validity. Rather, it indicates that the current conventional TCD endpoints were more sensitive to the common ordered demand than to the distinction between arithmetic and linguistic content. It also aligns with reports that conventional flow-velocity responses are not invariably scalable across cognitive paradigms (Intharakham, et al., 2022). Information-theory features may eventually add discriminant sensitivity, but they require a prespecified parameter strategy and independent validation.

### 4.3 Meaning of multimodal convergence

Behavioral and physiological measures changed with task level, and several pooled correlations linked slower or less accurate performance to altered TCD dynamics. Yet participant-specific behavioral slopes did not reliably track participant-specific physiological slopes. This distinction is important. The data support convergence at the experimental-manipulation level: higher levels change both performance and hemodynamic dynamics. They do not establish that an individual with twice the behavioral decline necessarily has twice the physiological change, nor do the correlations establish causal direction.

For experimental validation, this pattern is sufficient to show that task manipulation has observable consequences in two measurement modalities. For individualized cognitive-load estimation, however, larger samples, repeated sessions, stronger quality control, and models of blood pressure and CO₂ would be required. The present protocol is therefore better positioned as a group-level experimental framework than as a calibrated individual biomarker.

Measurement precision is increased by combining modalities. Behavioral measures provide direct evidence of achieved task difficulty, whereas physiological measures provide continuous and task-independent evidence of cerebral response. Neither modality alone is sufficient for calibrated individual cognitive-effort estimation: behavioral endpoints are influenced by prior ability, strategy, familiarity, and saturation, while TCD endpoints are influenced by systemic physiology, signal quality, blood pressure, and CO₂ regulation. A multimodal approach is therefore more defensible than inferring cognitive effort from either performance tests or physiological signals alone (Charles & Nixon, 2019).

### 4.4 Strengths and limitations

Strengths of this study include its within-subject design, the use of two cognitive domains and five ordered difficulty levels, software-controlled exposure and retention intervals, continuous bilateral TCD recording, and an auditable reanalysis that clearly distinguishes behavioral from physiological denominators and primary conventional endpoints from exploratory subjective and nonlinear measures. Several limitations should nevertheless be considered. The valid TCD samples were smaller and differed by domain, and only 18 participants contributed usable recordings for both paradigms. This reduced effective sample limits statistical power for detecting small domain-by-level, level-by-hemisphere, and three-way interaction effects; consequently, nonsignificant interactions should not be interpreted as definitive evidence that domain-specific or hemispheric effects are absent, because Type II error remains possible. This cautious interpretation is consistent with recent fTCD laterality work showing that hemispheric indices require explicit modeling choices, adequate signal quality, and sufficient statistical power before they can be interpreted as robust lateralized cognitive signatures (Thompson et al., 2023).

In addition, the fixed ascending presentation order prevented separation of task difficulty from elapsed time, habituation, learning, disengagement, and cumulative fatigue. No usable end-tidal CO₂ data were available, and systemic blood-pressure contributions were not included in the current endpoint models. NASA-TLX was collected only at the domain level, limiting its ability to assess the five-level gradient, and its interpretation requires consideration of the discrepancy between subjective workload and objective performance. Finally, arithmetic item-level responses were unavailable, and the information-theory analyses involved extensive parameter screening and should therefore be regarded as exploratory.

### 4.5 Future Work

Future studies will investigate whether graded levels of cognitive effort and hemispheric lateralization can be identified using cerebral blood-flow velocity measures alone. This work will compare conventional MCAv endpoints with statistical-complexity and entropy-based features. Validation should include prespecified feature sets, participant-level cross-validation, independent test samples, larger cohorts, counterbalanced task order, and simultaneous monitoring of blood pressure and end-tidal CO₂ to assess whether CBv-derived measures can support a reproducible physiological estimator of cognitive effort.

## 5. Conclusions

Five-level arithmetic and linguistic working-memory paradigms produced robust behavioral gradients and significant level-dependent changes in conventional TCD endpoints. The linguistic paradigm produced a steeper objective performance decline but reached an upper-level floor, while the arithmetic progression remained more gradual. The evidence supports orderly manipulation, but not psychometrically equidistant linguistic levels. Cerebral hemodynamic dynamics changed with level, but no robust domain-specific or hemispheric effects were detected. Behavioral and physiological measures therefore support a multimodal validation of the graded experimental manipulation, rather than separate arithmetic and linguistic neurovascular signatures. Future refinement should recalibrate the upper linguistic levels, standardize TCD quality-control and baseline rules, recover arithmetic item-level data, and evaluate the protocol in larger repeated-session samples with systemic covariates.

## Acknowledgments

This research was funded by ANID National Doctoral Scholarship N° 21221884 and the FONDECYT Regular Project 1241202.

## Appendix

## Appendix A1. Design and Operationalization of the Graded Arithmetic and Linguistic Working-Memory Tasks

### 1. Rationale and design principles

The arithmetic and linguistic tasks were developed to induce progressively increasing cognitive demands while allowing the simultaneous acquisition of behavioral and cerebral hemodynamic measures.

The tasks were not conventional n-back tasks because participants were not required to compare each stimulus with the stimulus presented (n) positions earlier. Instead, they were custom graded working-memory tasks inspired by experimental paradigms that manipulate processing difficulty, information retention, and response demands.

The design was informed by previous studies involving working-memory load, neurovascular coupling, transcranial Doppler monitoring, and cognitive-load theory. Attention was given to the limited processing and storage capacity of working memory and to the need to avoid excessive fatigue, frustration, or participant withdrawal.

The following general design principles were applied:

- Tasks had to be feasible for university students from scientific and humanities-related disciplines.
- Difficulty had to increase progressively without exceeding the expected cognitive capabilities of the target population.
- Each task had to include a retention interval between stimulus presentation and response.
- The total duration of each level had to remain approximately comparable across conditions.
- Rest periods had to be included to reduce cumulative fatigue.
- The interface had to minimize distraction and maintain attention on the stimulus.
- Behavioral performance and cerebral blood-flow velocity had to be recorded under standardized conditions.

The independent variables were cognitive domain and task level. The main dependent variables were behavioral accuracy, completion time, and baseline-normalized cerebral blood-flow velocity endpoints.

### 2. Arithmetic task design

#### 2.1 Arithmetic difficulty model

The arithmetic paradigm initially included addition, subtraction, multiplication, and division. Pilot testing indicated that the inclusion of multiple operation types increased variability and the risk of task abandonment. Therefore, the final paradigm was restricted to addition operations.

Arithmetic difficulty was manipulated through three components:

1. number of digits in each operand;
2. presence or absence of carrying;
3. number of operations included in each task.

The arithmetic difficulty score was defined as:

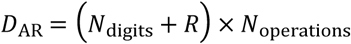

where:

- *N*_digits_ is the number of digits in the operands;
- *R* indicates whether carrying was required;
- *N*_operations_ is the number of additions included in the task.

**Table A1.** Arithmetic difficulty factors by level.

| Level | Number of digits | Carrying | Operations per task | Total difficulty |
| --- | --- | --- | --- | --- |
| AR-1 | 1 | 0 | 1 | 1 |
| AR-2 | 2 | 1 | 1 | 3 |
| AR-3 | 3 | 1 | 1 | 4 |
| AR-4 | 2 | 1 | 2 | 6 |
| AR-5 | 2 | 1 | 3 | 9 |

This operationalization increased either the numerical complexity, the need to maintain intermediate results, or the number of sequential operations.

#### 2.2 Arithmetic timing configuration

Each arithmetic task consisted of three sequential phases:

1. stimulus presentation;
2. retention interval;
3. response period.

The number of tasks and phase durations were adjusted to maintain a total duration of approximately 330 to 340 seconds per level. Comparability can be stated within each domain, but not between both.

**Table A2.**
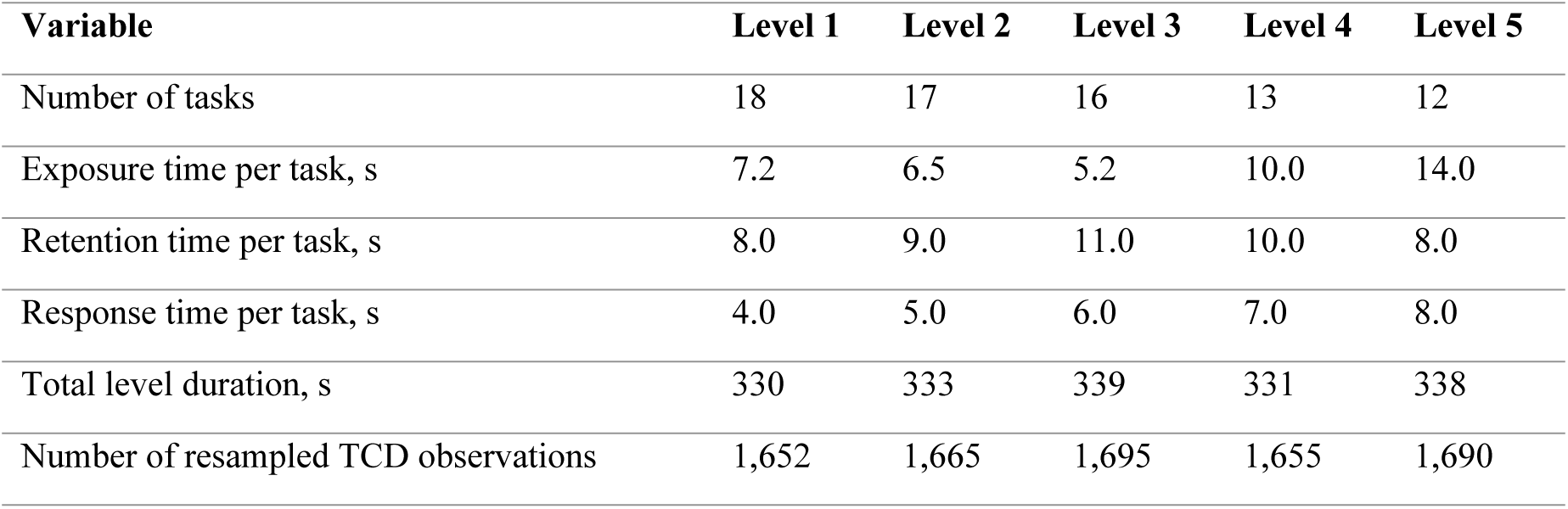
Arithmetic task configuration.

| Variable | Level 1 | Level 2 | Level 3 | Level 4 | Level 5 |
| --- | --- | --- | --- | --- | --- |
| Number of tasks | 18 | 17 | 16 | 13 | 12 |
| Exposure time per task, s | 7.2 | 6.5 | 5.2 | 10.0 | 14.0 |
| Retention time per task, s | 8.0 | 9.0 | 11.0 | 10.0 | 8.0 |
| Response time per task, s | 4.0 | 5.0 | 6.0 | 7.0 | 8.0 |
| Total level duration, s | 330 | 333 | 339 | 331 | 338 |
| Number of resampled TCD observations | 1,652 | 1,665 | 1,695 | 1,655 | 1,690 |

The average arithmetic level duration was approximately 334 seconds, corresponding to approximately 1,670 TCD observations after resampling.

#### 2.3 Arithmetic task sequence

For each arithmetic task:

1. the arithmetic stimulus was displayed;
2. the stimulus was removed;
3. a retention interval was presented;
4. the participant entered the result;
5. the next task began automatically.

The sequence was managed by experimental software to ensure standardized timing across participants.

### 3. Linguistic task design

#### 3.1 Linguistic task structure

The linguistic paradigm consisted of reading-comprehension tasks followed by a multiple-choice question. Participants had to retain and integrate information from the text before selecting a response.

Linguistic difficulty was manipulated using multiple textual and cognitive attributes. These attributes were combined into a study-specific difficulty index.

#### 3.2 Linguistic difficulty index

The linguistic difficulty score was defined as:

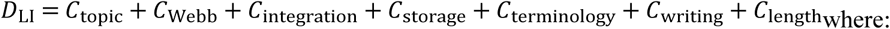

- *C*_topic_: topic complexity, scored from 1 to 5;
- *C*_Webb_: cognitive complexity based on Webb’s Depth of Knowledge, scored from 1 to 4;
- *C*_integration_: number of embedded or nested textual relationships;
- *C*_storage_: number of key concepts to be retained;
- *C*_terminology_: number of domain-specific terms;
- *C*_writing_: linguistic style and syntactic complexity;
- *C*_length_: number of words.

The linguistic difficulty index was developed specifically for this experiment. It was not intended as a previously validated psychometric scale, but as an operational tool for constructing ordered task conditions.

#### 3.3 Attribute scoring

##### Topic complexity

- 1: very low; 2: low; 3: moderate; 4: high; 5: very high.

##### Webb’s cognitive complexity

- 1: recall and reproduction; 2: skills and concepts; 3: strategic thinking; 4: extended thinking.

##### Storage demand

- 1: 1 to 3 key concepts; 2: 4 to 6 key concepts; 3: 7 or more key concepts.

##### Terminological repertoire

- 1: 1 to 3 topic-specific terms; 2: 4 to 6 topic-specific terms; 3: 7 or more topic-specific terms.

##### Writing complexity

- 1: simple informal writing; 2: intermediate or conversational writing; 3: formal and syntactically complex writing.

##### Text length

- 1: 40 to 80 words; 2: 81 to 120 words; 3: 121 to 150 words; 4: 151 to 200 words.

#### 3.4 Linguistic difficulty ranges

The following ranges were used to assign the texts to ordered levels:

##### Level Linguistic difficulty score

1. 3–4
2. 5–7
3. 8–11
4. 12–16
5. 17 or higher

These values defined ordered experimental conditions rather than psychometrically equidistant increments.

#### 3.5 Topics by level

The following topics were included:

- **Level 1:** principles of computer science and introduction to psychology;
- **Level 2:** computer science, environmental science, introductory microeconomics, introductory macroeconomics, and introductory statistics;
- **Level 3:** art history, world history, elementary biology, and elementary physics;
- **Level 4:** music theory, introductory calculus, intermediate chemistry, and English literature;
- **Level 5:** social impact of artificial intelligence.

Topic familiarity was expected to vary between participants and was therefore recognized as a potential source of additional variance.

#### 3.6 Linguistic timing configuration

The duration of each task was adjusted according to text length and complexity. Longer texts required greater exposure and response times. The number of tasks per level was reduced as task duration increased so that the total duration remained approximately comparable across levels.

**Table A3.**
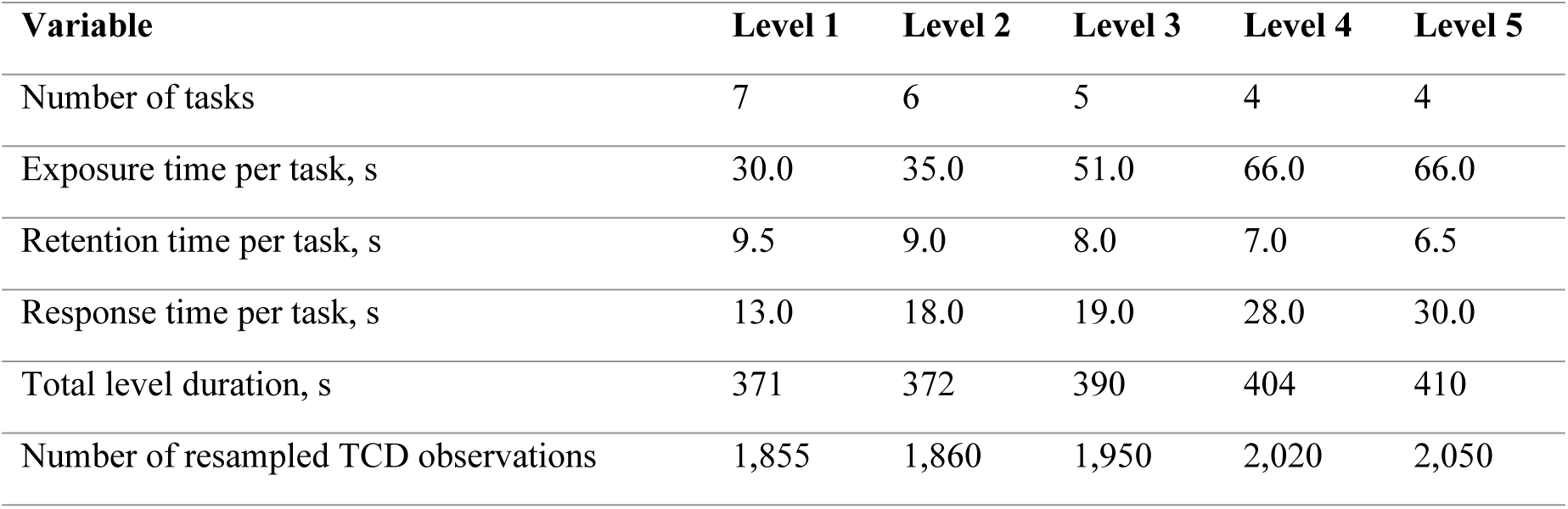
Linguistic task configuration.

| Variable | Level 1 | Level 2 | Level 3 | Level 4 | Level 5 |
| --- | --- | --- | --- | --- | --- |
| Number of tasks | 7 | 6 | 5 | 4 | 4 |
| Exposure time per task, s | 30.0 | 35.0 | 51.0 | 66.0 | 66.0 |
| Retention time per task, s | 9.5 | 9.0 | 8.0 | 7.0 | 6.5 |
| Response time per task, s | 13.0 | 18.0 | 19.0 | 28.0 | 30.0 |
| Total level duration, s | 371 | 372 | 390 | 404 | 410 |
| Number of resampled TCD observations | 1,855 | 1,860 | 1,950 | 2,020 | 2,050 |

The average linguistic level duration was approximately 390 seconds, corresponding to approximately 1,950 TCD observations after resampling. Comparability can be stated within each domain, but not between both.

#### 3.7 Linguistic task sequence

For each linguistic task:

1. a text passage was displayed;
2. the passage was removed;
3. a retention interval followed;
4. a multiple-choice question was presented;
5. the participant selected an answer;
6. the next task began automatically.

### 4. Experimental interface

The experimental software presented the stimuli using a visually simple interface. The display used a dark background with high-contrast text and included only the elements required for task completion.

The interface was designed to:

- minimize visual distractions;
- maintain central fixation;
- standardize exposure and retention times;
- record responses automatically;
- synchronize behavioral events with TCD acquisition;
- reduce experimenter intervention.

### 5. Baseline and duration matching

The 330s baseline was comparable to the arithmetic blocks, but shorter than several linguistic blocks. The baseline-normalized physiological endpoint was computed as shown in eq. (1). The same baseline file was used in the current analysis for both task domains.

### 6. Interpretation of the task levels

The five levels were designed as ordered conditions for increasing processing and retention demand. However, the levels should not be interpreted as equally spaced units of cognitive effort.

Behavioral validation subsequently showed:

- a progressive decline in arithmetic accuracy;
- a steeper decline in linguistic accuracy;
- increasing completion time in both domains;
- a nonlinear linguistic pattern with a floor region at the upper levels.

Accordingly, the levels are best described as experimentally ordered difficulty conditions rather than psychometrically equidistant categories.

### 7. Methodological considerations

Several factors should be considered when interpreting the task design:

1. Linguistic topic familiarity may have influenced performance independently of nominal difficulty.
2. The linguistic task showed evidence of a floor effect at levels 3 to 5.
3. If levels were always presented in ascending order, difficulty may be partially confounded with fatigue, habituation, or elapsed time.
4. The linguistic and arithmetic paradigms differed in stimulus structure, exposure time, and response format.
5. The difficulty indices were designed for experimental control and were not validated as standalone psychometric instruments.

### 8. Summary

The arithmetic and linguistic paradigms were designed to induce graded cognitive demand through progressively increasing requirements for processing, retention, integration, and response.

The arithmetic task manipulated numerical complexity and the number of sequential operations. The linguistic task manipulated textual, semantic, and memory-related attributes.

Although both paradigms produced ordered behavioral changes, the linguistic condition showed a steeper and nonlinear decline. Therefore, experimental manipulation should be interpreted as a multimodal graded design whose effectiveness was evaluated through behavioral and cerebral hemodynamic outcomes.

## Appendix A2. Gaussian Generalized Estimating Equation Models

To evaluate level-dependent changes in cerebral hemodynamic responses while accounting for the repeated-measures structure of the data, Gaussian Generalized Estimating Equation (GEE) models were fitted for each transcranial Doppler endpoint.

GEE models were selected because each participant contributed multiple correlated observations across cognitive domain, task level, and cerebral hemisphere. The method estimates population-averaged effects while providing robust standard errors in the presence of within-subject correlation.

A Gaussian distribution with an identity link function was specified because the physiological endpoints were continuous variables. An exchangeable working correlation structure was used, assuming a common correlation among repeated observations from the same participant. Participant identifier was defined as the clustering variable.

The general model specification for hemisphere-specific endpoints was:

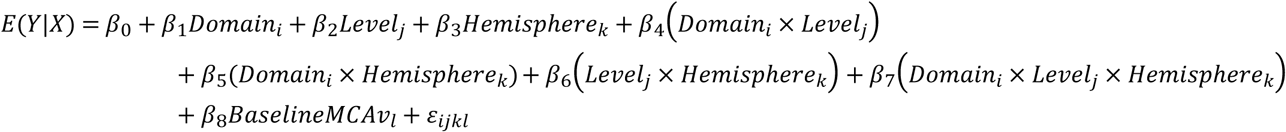

where:

- *Y_ijkl_* represents the TCD endpoint for participant (l), cognitive domain (i), task level (j), and hemisphere (k);
- *Domain* distinguishes arithmetic and linguistic tasks;
- *Level* represents the ordered task difficulty level from 1 to 5 and was modeled as a centered continuous variable;
- *Hemisphere* distinguishes left and right middle cerebral artery recordings;
- *BaselineMCAv* corresponds to the participant- and hemisphere-specific baseline mean cerebral blood-flow velocity;
- *β*_0_ is the model intercept;
- *β*_1_ to *β*_8_ are population-averaged regression coefficients;
- *ε* represents the residual error.

The corresponding model formula was:

*Y* ∼ *Domain* × *Level* × *Hemisphere* + *BaselineMCAv* For hemisphere-averaged endpoints, the hemisphere factor and its interaction terms were omitted:

*Y* ∼ *Domain* × *Level* + *BaselineMCAv* The primary physiological endpoint was the baseline-normalized percentage change in mean middle cerebral artery blood-flow velocity included in eq. 1. Additional continuous TCD endpoints analyzed using the same Gaussian GEE framework included: peak percentage change in MCAv; net area under the curve; positive area under the curve; time to peak; slope to peak; early 30-second slope; final 10-second residual change; peak-to-final change.

The principal coefficient of interest was the main effect of task level, which quantified the average change in each physiological endpoint for each one-level increase in task difficulty. Interaction terms were used to test whether this level-related change differed between arithmetic and linguistic domains or between hemispheres.

For the primary endpoint, the model identified a significant negative level effect on baseline-normalized mean MCAv: *β* = −1.084, *SE* = 0.257, *p* = 2.40 × 10^−5^, with a 95% confidence interval from (−1.59%) to (−0.58%). This result indicates that each one-level increase in task difficulty was associated, on average, with an additional decrease of approximately 1.08 percentage points in mean MCAv relative to baseline.

No statistically significant domain-by-level interaction was observed for mean MCAv: *β* = −0.193, *p* = 0.604, and the three-way interaction between domain, level, and hemisphere was also not significant: *β* = −0.489, *p* = 0.302.

Therefore, the GEE results support a robust population-level effect of graded task difficulty on cerebral hemodynamic responses, but they do not provide evidence of distinct domain-specific or hemisphere-specific progression patterns.

Robust standard errors were used for inference. Model coefficients were interpreted as population-averaged associations rather than subject-specific trajectories. Since the available dataset did not include usable end-tidal carbon dioxide measurements, no EtCO_2_ adjustment was performed, and no claim of CO_2_-controlled neurovascular response was made.

The GEE analyses were considered appropriate for the primary population-level inference because they retained all valid repeated observations and accommodated within-subject dependence without requiring successful estimation of subject-specific random effects. However, the interpretation of the physiological results remained conditional on the finalized signal-quality inclusion criteria and baseline definition.

## Appendix A3. Direction and precision of significant TCD level effects forest plot

**Figure A3.1.**
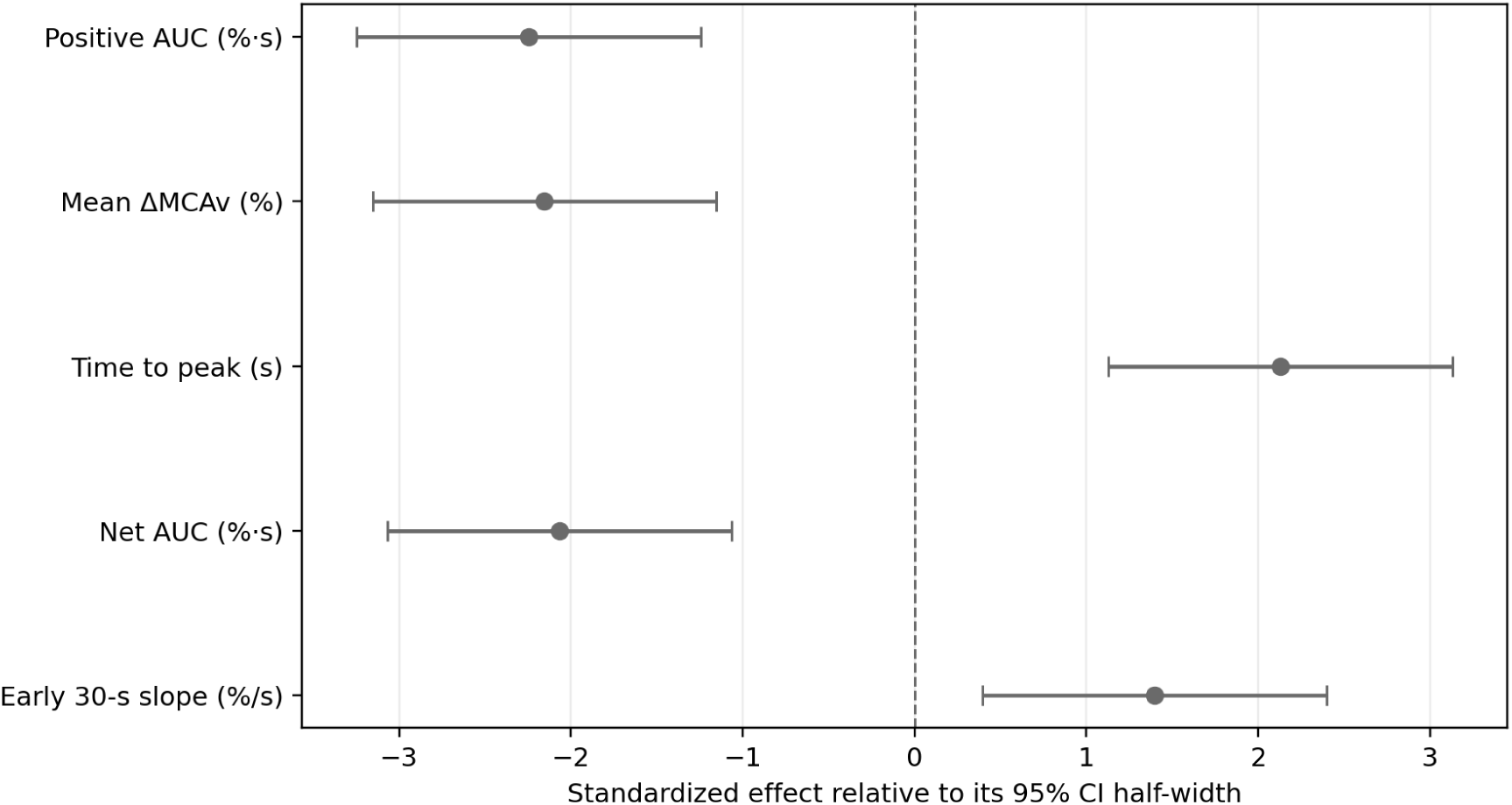
Direction and precision of significant TCD level effects. The standardized display allows endpoints with different units to be shown together; exact coefficients and confidence intervals are reported in Table 5.

